# Evolutionary constraints improve protein large language model predictions for protein stability, binding regions and epistasis

**DOI:** 10.64898/2026.05.22.726784

**Authors:** Konstantina Tzavella, Catharina Olsen, Wim Vranken

**Affiliations:** Department of Medical Physics, Institut Jules Bordet, Hôpital Universitaire de Bruxelles (H.U.B), Université Libre de Bruxelles (ULB), Meylemeersch 90, Brussels, 1070, Belgium; Interuniversity Institute of Bioinformatics (IB2), ULB-VUB, Brussels, Belgium; Brussels Interuniversity Genomics High Throughput Core (BRIGHTcore), Vrije Universiteit Brussel (VUB), Université Libre de Bruxelles (ULB), Brussels, Belgium; Clinical Sciences, Research Group Reproduction and Genetics, Centre for Medical Genetics, Vrije Universiteit Brussel (VUB), Universitair Ziekenhuis Brussel (UZ Brussel), Laarbeeklaan 101, Brussels, 1090, Belgium; Structural Biology Brussels, Vrije Universiteit Brussel, Brussels, Belgium; Chemistry Department, Vrije Universiteit Brussel, Brussels, Belgium; AI Lab, Vrije Universtiteit Brussel, Pleinlaan 2, Brussels 1050, Belgium

**Keywords:** protein stability prediction, intragenic epistasis, mutation co-occurrence effect prediction, binding site prediction, protein Language Models

## Abstract

Our understanding of protein function and evolution is largely based on the relationship between amino acid sequence and overall fold, now effectively captured by computational models. Yet predicting how mutations—shaped by epistasis—alter protein behavior, especially in dynamic or structurally ambiguous regions, remains difficult. Here we present D2D, which combines a self-supervised protein language model with protein-specific evolutionary information to predict mutational effects using little to no task-specific labeled data. D2D captures long-range epistatic interactions, accurately predicts single and higher-order mutation effects on protein thermostability and binding, without being trained on the task. When fine-tuned, D2D outperforms state-of-the-art methods on latent driver cancer mutations and co-occurring proliferation-enhancing mutations across independent experimental studies. Unlike most existing approaches, D2D avoids biases linked to solvent accessibility or to multiple sequence alignment depth and quality, making it particularly effective for disordered or surface binding regions where structure-based predictors typically falter. Overall, D2D provides a general framework for modeling mutational effects in proteins with limited experimental or structural information.

## Introduction

Proteins are the molecular machines of all living systems, performing essential cellular functions from structural support and enzymatic activity to molecular transport. Their function is often determined and constrained by the three-dimensional structure they adopt. In such a structure, amino acid residues interact so that changes at one site may require compensatory adjustments elsewhere to preserve packing, electrostatics, or flexibility [1-4]. These context-dependent mutational effects, known as intragenic epistasis, are pervasive in protein evolution [5], [7-9], reverse genetic drift in small populations [10], [11], restore protein activity and influence drug resistance in cancer therapy [6, 12, 41, 42]. For example, secondary mutations in BRCA2 can counteract primary mutations that are initially hypersensitive to chemotherapy drugs like cisplatin. By restoring the wild-type reading frame and protein structure of BRCA2, these secondary mutations ultimately drive the development of drug resistance.

Due to the complexity of proteins and amino acid interactions, quantitatively predicting or modelling the outcomes of intragenic epistasis remains difficult. The intricate behavior of proteins, the ‘protein phenotype’, is typically categorised by stability, which is highly relevant for well-folded protein domains such as enzymes, and functionality, which is typically closely related to conserved amino acids at functional sites or binding interfaces and motifs. Disrupting stability will however often also affect functionality, and separating these effects is not trivial, especially when multiple mutations are present.

Traditionally, multiple sequence alignments (MSAs), which capture evolutionary information (EI) on protein families, have been the established framework for inferring constraints on protein phenotype computationally [13-15]. Although MSAs provide explicit estimates of evolutionary constraints and residue coupling, their utility can be limited by alignment depth, family diversity, and computational demands, particularly when modeling higher-order dependencies such as interactions between multiple residues that mediate long-range allosteric effects. Nevertheless, most current computational predictors only evaluate the effect of single mutations or approximate multi-mutant outcomes under the assumption their contributions are additive [16-17].

More recently, protein Language Models (pLMs) have emerged as powerful tools in predicting tertiary structures [18], intrinsic disorder [19], single-cell annotations [20] and protein-protein interactions [21], even validating biological hypotheses [22]. Trained in an alignment-free manner on massive sequence datasets, they produce rich but extremely high-dimensional embeddings, typically ∼1000 per amino acid residue. This large embedding space poses a practical challenge for downstream models, particularly in data-limited settings, where it increases the risk of overfitting with many more features than samples. Moreover, unrealistic predictions of protein isoform structures have revealed that pLMs do not truly grasp the biophysics of proteins. Instead, they rely on statistical coevolutionary patterns, learned agnostically from many thousands of protein families [23].

The question that then poses itself is how and whether specific evolutionary information for a protein family, as captured in MSAs, can still add to the statistical, general pLM representations. In our D2D approach, we explore this question by combining the strengths of both: for each protein family, we constrain the general statistical representations learned by pLMs with protein-specific evolutionary information derived from MSAs. By modeling site-specific variation in embedding space using a Gaussian Mixture Model (GMM), we approximate the distribution of evolutionarily permissible states at each position. Mutational effects are then quantified as deviations from this distribution, enabling detection of substitutions that are statistically plausible in a global sense but incompatible with the local evolutionary constraints of the protein under study.

We have previously demonstrated the model’s effectiveness in predicting single cancer driver mutations [24] and now propose its general applicability to other biologically relevant tasks involving protein (mis-)behavior, such as protein stability prediction and binding-site identification. This extended sequence-based framework (D2D) accurately identifies binding sites, including those located within intrinsically disordered regions, even under limited or no labeled data. Furthermore, it successfully detects novel, latent co-occurring mutations whose combined effects are associated with poorer treatment outcomes, capturing the non-linear interactions that contribute to cancer progression.

**Figure 1:**
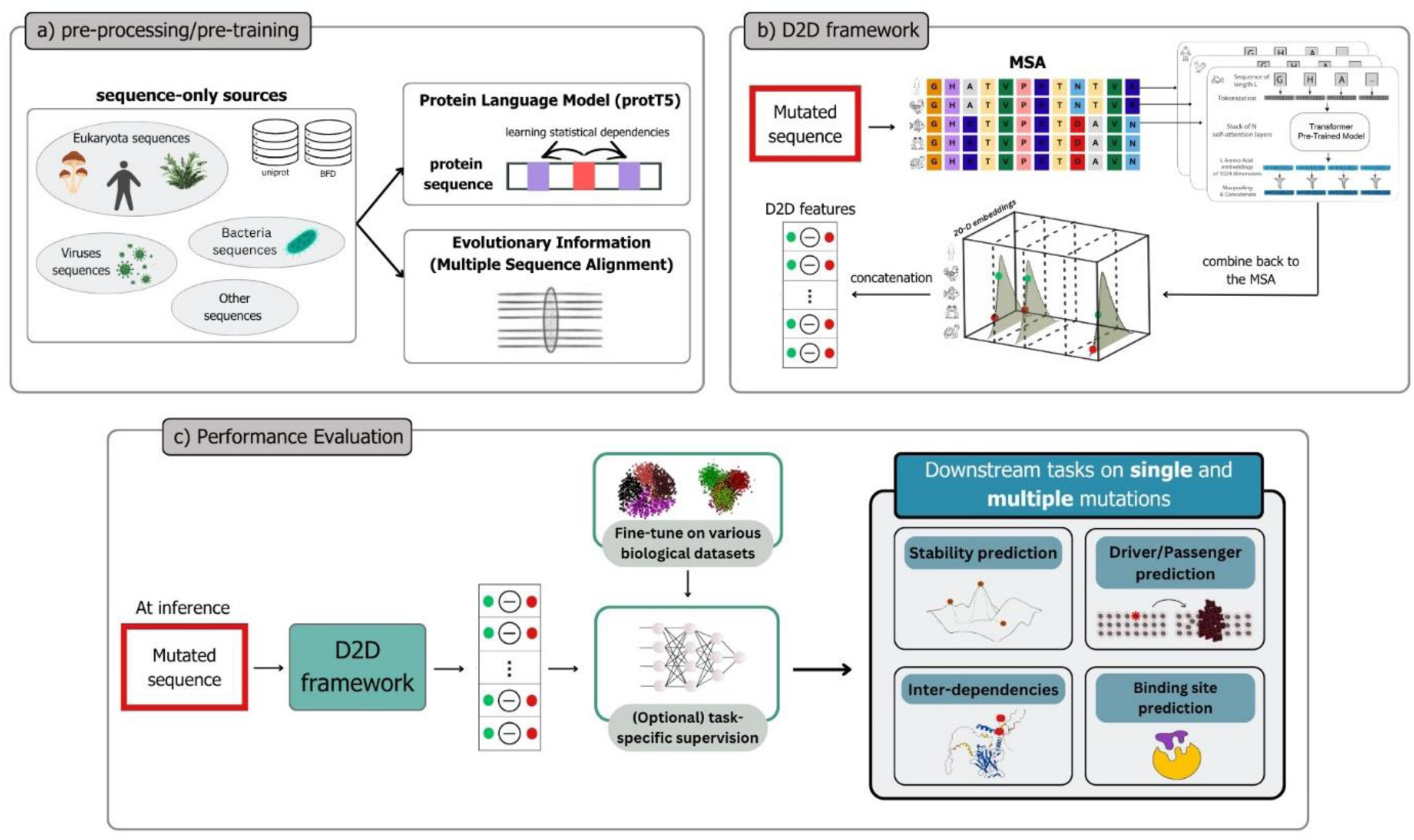
D2D pipeline. (a) A protein Language Model pre-trained on millions of protein sequences captures the statistical dependencies between amino acids. The Multiple Sequence Alignments tool mmseqs2 uses the UniProt and BFD databases and captures the protein-specific distribution of the protein site. (b) Core D2D framework. Each amino acid of the MSA is mapped to a selected 20-D pLM representation vector. The pLM vectors are combined back to the MSA and a Gaussian Mixture Model (GMM) is fitted at each position across species. The log-probability of the mutant representation is compared to that of the wild type, and the differences are concatenated to quantify mutational effects across the sequence. A one-dimensional GMM is shown for illustration. (c) The resulting features can be used in an unsupervised manner (e.g. averaging) or as input to supervised models for downstream tasks. Outputs include prediction and confidence scores and enable identification of short- and long-range amino acid interactions.

## Results

To assess the broad effectiveness of combining pLM and EI for both single and multiple mutations, we focused on three key protein phenotypes: (i) protein stability, (ii) binding site prediction with emphasis on Linear Interacting Peptides (LIPs) that mediate protein-protein interactions, and (iii) the impact of secondary, synergistic mutations on tumor growth.

### D2D demonstrates strong performance in protein stability and compensatory mutations detection across solvent accessibility levels

Amino acid mutations that alter the thermodynamic stability of proteins are associated with various diseases. In cancer, for example, such mutations can cause tumor suppressor proteins to unfold, or can conversely stabilize oncogenic proteins. To evaluate the performance of our model on thermodynamic stability data, we used the Mega-scale dataset [25], which quantifies the effects of single amino acid substitutions on the free energy difference between folded and unfolded states (ΔΔG). We benchmarked on 291 776 mutations, including 162 725 single and 129 051 double mutations, across 166 domains.

We benchmarked D2D against state-of-the-art (SOTA) predictors GEMME [26], mSCM [27], MSA_transformer [28] and ESM-1v [29] that leverage evolutionary, structural, protein language model and evolutionary/protein language model information respectively. The unsupervised D2D approach, which is agnostic of thermodynamic stability information, already achieves Spearman correlation scores comparable to SOTA methods, significantly outperforming the well-established, specifically trained mCSM predictor (Wilcoxon test p-value=0.003) and MSA_Transformer (p-value=0). Because it captures the statistics of the variation of pLM embeddings, D2D can provide a confidence score for each prediction based on the quality of the Gaussian fit to the underlying MSA data. When restricting the analysis to the 24 studies with average confidence scores at least 60%, the Spearman correlation increased to 0.58, outperforming all other predictors (Fig.2A, left panel). Confidence scores above 70% were excluded from further analysis due to insufficient data points. In agreement with the conclusions of [24], D2D features demonstrate significantly superior performance compared to those derived from the ProtT5 pre-trained model in a supervised setting with cross-validation (Fig.2A, right panel).

**Figure 2:**
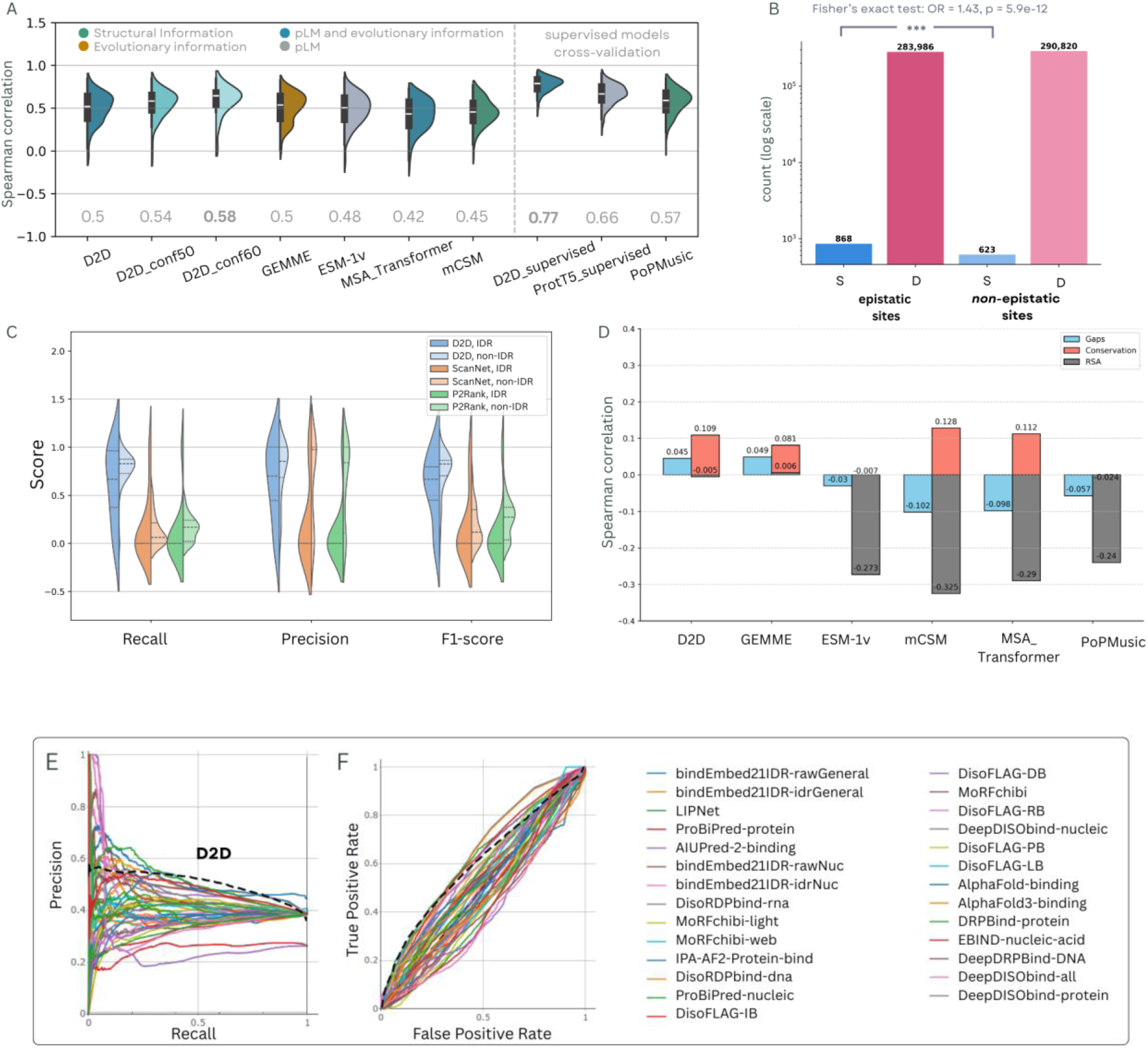
(a) D2D predictions compared to SOTA predictions for the mega-scale dataset of experimental ΔΔG values upon single and double mutations. D2D_conf50 and D2D_conf60 represent predictions made at 50% and 60% confidence thresholds, respectively. (b) Epistatic sites were defined as positions where the D2D signal exceeded one-tenth of its maximum value. Compensatory mutations (secondary mutations that alleviate the negative effects of a single mutation) occur more frequently (Odds Ratio ≈ 1.43, p ≈ 6×10−12, Fisher test) on epistatic sites than on non-epistatic sites, (c) Comparison of D2D-derived LIP predictions with ScanNet and P2Rank, (d) Spearman correlations between the performance across 166 mega-scale studies and the corresponding average Gaps, Conservation, and RSA values per study. A negative Spearman correlation (e.g., for RSA) indicates that predictors perform worse for exposed residues (i.e., residues with high RSA), (e) Performance of the unsupervised D2D model on the CAID-3 Binding-IDR dataset, achieving an average AUC of 0.64, average APS of 0.50, and F1-max of 0.56 (a comprehensive overview of the performance of all predictors is provided in Supplementary Table 1).

A major challenge the stability predictors face is their inherent biases built into their design. For evolutionary predictors, poor-quality MSAs can reduce the reliability of the conservation score, potentially affecting the overall pathogenicity prediction. Structure-based predictors exhibit a different bias, tending to classify mutations buried (low Relative Solvent Accessibility (RSA)) in the protein core as pathogenic [32]. To investigate these tendencies, we evaluated the dependence of each predictor on key parameters: the proportion of gaps at the mutation site in the MSA, the conservation level at the mutated position, and the residue’s RSA.

Consistent with previous observations [33], [34] the performance of the two structure-based predictors mCSM and PoPMusic deteriorated with the increase of RSA (Fig.2D). In other words, structure predictors struggle to correctly estimate the impact of mutations in exposed residues, as residues are not packed together, making change in amino acid side chains more difficult to assess. Interestingly, the sequence-based, pLM predictors MSA_Transformer and ESM-1v also appeared to be impacted by the RSA of the residue. In contrast, evolutionary models D2D and GEMME remain robust across varying RSA levels. This discrepancy may stem from two factors: (i) structure-based methods often rely on static structural representations that inadequately capture the dynamic nature of exposed regions, and (ii) mutations at exposed sites may primarily impact functional properties -such as binding or catalysis- rather than stability, which most ΔΔG predictors are designed to assess. Regarding the MSA conservation, we see a minimal impact on the performance, especially for mCSM, with almost all predictors performing better when conservation is higher. The presence of gaps in the MSA does not have any effect.

To investigate D2D’s ability to deal with multiple mutations we focused on the compensatory mutations, where the negative effects of one mutation are alleviated by epistatic interaction with a second mutation at another site. Although not fully understood, this phenomenon plays a critical role in important biological processes like antibiotic resistance [40] and cancer therapy [42]. In tumors, for instance, mutations in sites targeted by drugs can directly or indirectly interfere with their binding, leading to drug resistance [41].

We hypothesized that by constraining the protein-agnostic coevolutionary statistics with protein-specific evolutionary information, we can uncover relationships between mutations and protein sites that are prone to interact. To do so, we first defined sites as epistatic when their D2D signal was greater than 10% of the global maximum D2D signal across all sites. From the mega-scale dataset, we then selected single mutations that are destabilizing (ΔΔG < −2) but become stabilizing (ΔΔG > 1) upon introduction of a second mutation, following the stabilizing threshold proposed in [25]. The double stable mutants on epistatic sites were 299 compared to 198 on non-epistatic sites. In contrast, destabilizing mutations were more frequently observed at non-epistatic sites than at epistatic ones (Fig.2B). Overall, double mutations that restore stability are enriched 1.4-fold at epistatic sites relative to non-epistatic sites (Odds Ratio ≈ 1.43, p ≈ 6×10−12, Fisher test) supporting our hypothesis.

### Binding sites prediction in ordered and disordered regions

Linear Interacting Peptides (LIPs) are functional motifs involved in binding and intermolecular interactions, mostly found in intrinsically disordered regions. In these poorly structured regions, sequence conservation is typically low, making it difficult to assess the impact of mutations, even though their functional impact can be high. We therefore sought to evaluate whether the combination of pLMs and EI features works in this challenging case.

We compared LIPs from the mega-scale collection, annotated in the MobiDB database, with interaction sites predicted using our approach as described in Methods. These results were benchmarked against two state-of-the-art predictors: P2Rank [30] and ScanNet [31]. P2Rank is a Machine Learning method trained to predict ligand binding sites from protein structures. ScanNet, is a structure-based predictor that uses geometric deep learning to learn spatio-chemical patterns of binding sites directly from 3D structures, also specifically trained to predict binding sites.

We hypothesized that mutations with minimal impact on protein stability but strong evolutionary constraint are enriched in functional sites. To investigate this, we combined experimental stability measurements (ΔΔG) with evolutionary scores to construct mutational landscapes capturing low stability impact and high evolutionary effect. We evaluated performance separately for intrinsically disordered regions (IDRs) and ordered regions. As summarized in Fig.2C, our method, D2D, outperforms both P2Rank and ScanNet for binding sites in both IDRs and ordered regions, although all predictors perform better on ordered regions compared to IDRs.

To further validate our approach, we tested D2D on the CAID-3 Challenge binding-IDR dataset, comprising 52 sequences, and compared its performance to the other algorithms that participated in the challenge (Fig.2E), confirming its superior predictive ability in disordered regions. The CAID evaluation framework currently permits only CPU-based execution under constrained computational resources. As discussed in the Computational Requirements section, these limitations prevent on-the-fly benchmarking of D2D within the CAID competition due to the GPU requirements associated with ProtT5 inference.

### Latent driver mutations discovery and their effect on tumor growth

As the D2D approach is relevant for single amino acid driver versus passenger classification in cancer [24], we further explored its performance on genes where double mutations increase oncogenic activity. Findings in [38] indicate that combining frequent driver mutations with rare or weaker, latent, mutations in the same gene can significantly enhance tumor progression and affect treatment outcomes. To investigate this, we used the mutations reported in that study and created a negative (control) set from gnomAD [39] for the same genes, as described in Methods. In total 25 992 pathogenic and 22 743 benign mutations across 18 genes were tested. Because of the class imbalance between the latent driver positions and gnomAD variants, the balanced accuracy metric was used to assess the minority class performance. D2D and GEMME results are shown on Fig.3A (left panel) with average weighted accuracy 0.63 for D2D and 0.44 for GEMME (Wilcoxon signed-rank test p-value=0.006). Consistently across all evaluated genes, D2D outperforms GEMME, as further illustrated in Fig. 3B.

**Figure 3:**
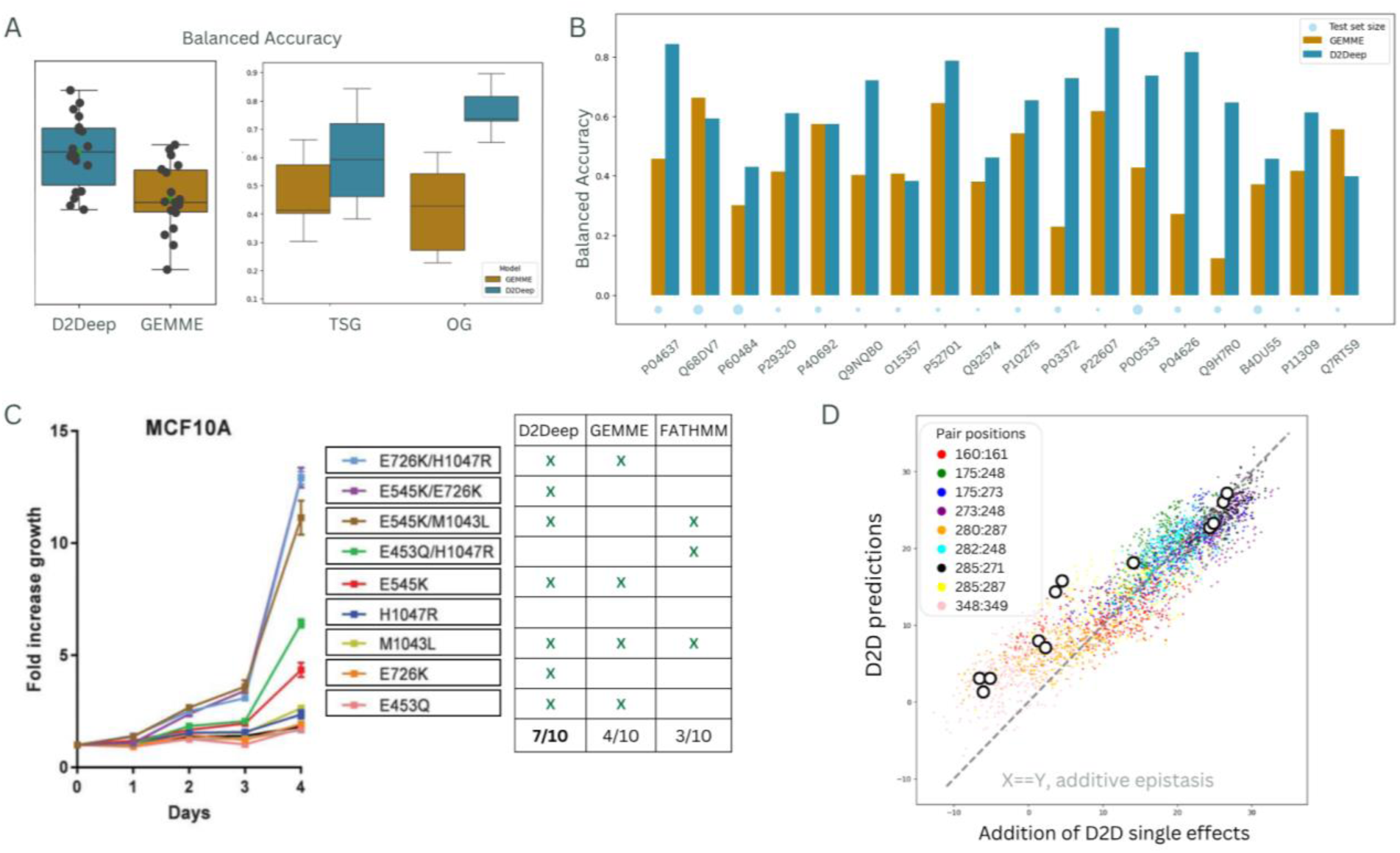
(a) Balanced accuracy for D2Deep and GEMME predictors for latent mutations [38] and gnomAD control mutations (left plane) - D2Deep demonstrates better results for Oncogenes (OG) compared to Tumor Suppressor Gene (TSG) (right plane), (b) Detailed performance of latent mutations and control mutations on 18 genes, (c) D2Deep, GEMME, and FATHMM-cancer mutation effect prediction for PIK3CA single and double mutants, (d) Comparison of additive vs. epistatic effects for TP53 double mutations using raw D2Deep predictions (pre-sigmoid). Mutation pairs were assessed across latent residue positions reported by [165], evaluating all amino acid combinations. The x-axis shows the sum of individual mutation effects, while the y-axis shows the corresponding D2Deep prediction for the double mutant. ‘O’ marks highlight the mutated amino acid combinations with known ground truth from the experimental study.

From Fig.3A (right panel) we also conclude that D2D performs better for Oncogenes (OG) compared to Tumor Suppressor Gene (TSG). This indicates that D2D is more sensitive in identifying the activation of OG (through gain-of-function mutations) than the inactivation of TSG (loss-of-function mutations), both of which are key processes in cancer development

We then focused on driver mutation pairs of the well-known TP53 gene, by comparing the simple addition of individual predictions with the epistatic dual mutant prediction, across all suggested epistatic positions suggested in [38] and for all amino acid combinations. The rationale underlying this analysis is that co-occurring mutations within the same gene may exhibit additive or cooperative effects. When the predicted effect of a double mutant equals the sum of the individual mutation effects, this indicates a lack of functional interaction. In contrast, when the phenotype of a double mutation cannot be explained by the additive effects of the individual single mutants, indicates epistatic links. When the combined mutations produce a phenotype that is more detrimental than expected, the interaction is classified as negative epistasis. In contrast, if the double mutant displays a milder phenotype than predicted, the interaction is considered positive.

The comparison between additive and epistatic predictions is shown in Fig. 3D, where the x-axis represents additive effects and the y-axis represents D2D epistatic predictions. The ‘O’ marks highlight the mutated amino acid combinations with known ground-truth from [38]. The majority are above the diagonal and so align with the study’s conclusions regarding negative epistatic contributions. The points with high prediction values cluster to the diagonal, suggesting that epistasis is not required to amplify their pathogenicity. This pattern is consistent in all positions with colored filled dots, where we identify mutation pairs for which individual variants are predicted to be benign (effect < 0), yet their combined epistatic prediction is pathogenic. This is observed, for instance, at the positions 248:249 and 285:287 that predominantly show negative epistasis effects. On the contrary, pairs such as 273:248 and 285:271 display positive epistatic interactions (Supplementary Fig. 5). We note here that for positions 348:349, a stop codon mutation is proposed in [38]. As shown on the last panel of Supplementary Fig. 5, D2D predictions for the remaining amino acid substitutions appear to have a lower impact on the protein’s function (predictions below zero).

Finally, to further illustrate the epistatic signal captured by D2D and its relevance on actionable clinical analysis, we selected PIK3CA, the most frequently mutated oncogene in human cancers. It has hotspot single–amino acid mutations commonly occurring in the helical domain (E542K, E545K) and kinase domain (H1047R). Clinical trials [35, 36] have shown that some patients with particular double PIK3CA mutations experience prolonged benefit from alpelisib, suggesting that such double mutations may serve as a biomarker for enhanced response to PI3K inhibitors. This prompted researchers to conduct an in-depth analysis of the prevalence and biological significance of these mutations in relation to PI3Kα inhibitor sensitivity. In [37], a growth proliferation time course study of PIK3CA mutant MCF10A cells, serum-starved over four days, revealed that cells with double mutations displayed increased proliferation compared to single mutants. We compared these experimental findings with the predictions from D2Deep, GEMME, and FATHMM-cancer models. D2Deep accurately predicted the increase of mutation effect in correct order for seven out of ten mutation effects, outperforming GEMME (four correct predictions) and FATHMM-cancer (three correct predictions) (Fig.3C).

## Discussion

Protein Language Models (pLMs) have shown remarkable potential for addressing diverse problems in computational biology. However, their general-purpose representations can struggle to capture specialized biological knowledge that depends on specific evolutionary or structural contexts. D2D addresses this limitation by fine-tuning the pLM-derived embeddings with explicit evolutionary information obtained from sequence alignments.

Here, we present a proof of concept of this approach by evaluating the predictive capability of D2D across several biological tasks, including protein stability and binding. In the context of stability prediction, D2D shows reduced bias with respect to residue accessibility compared with many current state-of-the-art approaches. In particular, the model is able to capture stability effects arising from mutations located on the protein surface that are epistatically coupled with residues in the structural core. This suggests that combining pLM representations with evolutionary constraints can help capture long-range dependencies that influence stability.

Interestingly, D2D also performs well on binding-related tasks despite not being explicitly trained on protein–protein interaction data. We interpret this as evidence that the relative contribution of the pLM and EI components shifts with the task. In stability prediction, columns of the MSA carry strong signal because structurally important residues tend to be highly conserved. In binding, however, residues that are critical for interaction but do not strongly affect stability can remain weakly conserved, limiting the information available from MSA-derived statistics alone. Here, the pLM component appears to provide complementary information: because masked language modeling predicts residues from sequence context, substitutions — even in flexible or poorly conserved regions — can produce large embedding perturbations that flag a functionally relevant change.

This finding suggests that models such as D2D could potentially be extended to large-scale analyses of intergenic epistasis and protein–protein interaction prediction. To achieve this, future extensions would need to incorporate representations capable of modeling inter-chain dependencies. One promising direction would involve replacing the current sequence embeddings with complex-aware representations derived from models trained on protein–protein complexes. In such a framework, the current approach of fitting one Gaussian Mixture Model (GMM) per residue position based solely on a protein’s multiple sequence alignment (MSA) could be generalized. Instead, GMMs could be trained on embeddings representing co-evolved residue pairs or conserved interface motifs. D2D could then compute two complementary quantities: intra-chain distances, measuring how well a mutant residue fits within the evolutionary distribution of its own protein, and inter-chain distances, reflecting how compatible that mutation is with the interface features of its binding partner.

Another strength of D2D is its robustness in data-limited settings. The model performs well even with weak supervision and small labeled datasets. Beyond single mutations, applying D2D to co-occurring variants provides a potential framework for analyzing mutational landscapes in cancer and may help identify functionally relevant mutation combinations for improved patient stratification.

Overall, our results demonstrate that refining pLM representations with evolutionary constraints within a probabilistic framework enables the analysis of mutational effects across diverse biological contexts with minimal supervision. Future work incorporating complex-aware representations and explicit modeling of co-evolving interfaces may further extend this approach to predicting interaction-specific epistasis and large-scale protein-protein interaction networks.

## Methods and Materials

### D2D Architecture and extension

This work builds on the method introduced in [24], in which a supervised D2D framework was developed for cancer driver mutation prediction. Briefly, for each mutated protein sequence, a genetic database search is performed to construct a multiple sequence alignment (MSA) across diverse species. Each amino acid in the MSA is then mapped to a 20-dimensional protein Language Model (pLM) representation vector, which is selected from the available embeddings by maxpooling. We reduced the feature dimensions using max-pooling with a kernel size of 50 and a stride of 50, resulting in 20-dimensional vectors (20-D) for each amino acid. Given 20 x 50 =1000, dimensions beyond this range (i.e., the final 24) were excluded from the analysis. Retaining these samples would require zero-padding the signal to the next multiple of 50; however, we opted not to do so, as padding introduces artificial values that could bias the pooled maxima. Since the omitted portion represents less than 2.3% of the total input length, its impact on the resulting features is negligible.

The 20-D representations are integrated back into the MSA, and a Gaussian mixture model (GMM) is fitted to the representations at each alignment position across species. For a given mutation, the GMM log-probability of the mutant representation is subtracted from that of the corresponding wild-type representation (Supplementary Fig. 3). The distributions of MSA depths are shown in Supplementary Fig. 1. The resulting differences are concatenated across all sequence positions to capture the global effect of the mutation that is either fed to a supervised learning model or averaged for the unsupervised extension of D2D. For the supervised setting, 5-fold cross-validation was employed separately for each protein, ensuring that mutations in the training and test sets were non-overlapping.

### Thermodynamics protein stability dataset

We leveraged a mega-scale dataset [25] that quantifies changes in thermodynamic stability (ΔΔG) between folded and unfolded states upon the introduction of single and double substitutions under standardized conditions. The dataset was generated using a high-throughput cDNA proteolysis assay in a cell-free in vitro translation system derived from Escherichia coli and comprises 331 natural protein domains and 148 designed proteins, each up to 72 amino acids in length, across a diverse set of organisms, measured under identical conditions.

To ensure data reliability, we excluded the domains with low-confidence ΔΔG measurements, as well as mutated structures that could not be mapped to a UniProt reference sequence. This resulted in 291 776 mutations, including 162 725 single and 129 051 double mutations, across 166 domains. Following the definition used in Tsuboyama et al [25], mutations with ΔΔG > 1 kcal/mol were classified as stabilizing. For this dataset, D2D predictions were generated in an unsupervised manner by averaging the corresponding D2D feature representations.

### MobiDB Linear Interacting Peptides sites and CAID-3 Challenge

Functional sites were annotated using MobiDB, a database that compiles disorder annotations for 245.5 million proteins, integrating experimentally validated and curated information on intrinsically disordered regions (IDRs) and Linear Interacting Peptides (LIPs). Protein domains from the mega-scale dataset were mapped to their corresponding UniProt sequences, accounting for cases where experimental constructs included truncated or extended regions. Disorder and LIP annotations were then retrieved for the mapped regions. We retained mutations with small stability effects (−2 < ΔΔG < 1 kcal/mol) and low D2D scores (below the 60th percentile; Supplementary Fig. 2). The D2D predictions were computed in an unsupervised way, by averaging the D2D features. For the functional annotation, we only considered high-confidence evidence from MobiDB, including experimentally supported annotations (Curated and Derived) and the highest-confidence disorder predictions, as detailed in Supplementary Table 2.

### Latent mutations analysis

To define an appropriate negative control set, we leveraged population variation data from the Genome Aggregation Database (gnomAD) database (https://gnomad.broadinstitute.org/). For each gene of interest - identified as latent candidates in [38] - we retrieved common variants with allele frequency greater than 0.5%, under the assumption that such variants are more likely to be tolerated. We then generated double mutations combining these common variants within each gene, producing a set of putatively benign mutation pairs (gnomAD_latent_pairs). In total 25 992 pathogenic and 22 743 benign mutations were tested in 18 genes that contained both latent driver pairs and corresponding gnomAD-derived variants (latent_driver_pairs). Predictions for this dataset were generated using the supervised D2Deep framework, as described in [24]. For the epistatic analysis, we used raw D2D prediction scores prior to sigmoid transformation. This choice preserves the full dynamic range of the model outputs, enabling direct comparison of absolute effect sizes between additive and epistatic predictions.

## Data Availability

We provide the D2D features calculated for the mega-scale and CAID_binding_idr datasets, the latent_driver_pairs and the gnomAD_latent_pairs in the Zenodo repository: https://doi.org/10.5281/zenodo.20309165

## Computational Requirements

The per-residue pLM embeddings and Gaussian Mixture Model fitting were performed on a TU102 GPU (GeForce RTX 2080Ti) equipped with 11GB of GDDR6 memory. As an alternative to on-the-fly embedding generation, precomputed embeddings from the UniProt database could be used. However, at the time of writing, embeddings are only available for reviewed sequences, while multiple sequence alignments may also include unreviewed entries. The framework requires Python 3.10 or later together with the following packages: PyTorch, Transformers, EVcouplings, scikit-learn, and Biopython. GPU acceleration is strongly recommended for ProtT5 inference.

## Code Availability

The D2D code, together with instructions for reproducing the results and the associated software and hardware requirements, is available at: https://github.com/KonstantinaT/D2D

## Funding

Vrije Universiteit Brussel Research Council under the Interdisciplinary Research Program TumorScope [IRP20 to K.T.]; European Union’s Horizon 2020 research and innovation programme under grant agreement No 101016834 (HosmartAI) to K.T., W.V.; Research Foundation Flanders (FWO) International Research Infrastructure [I000323N to W.V.].

## Conflict of interest/Competing interests

Not applicable

## Ethics approval

Not applicable

## Acknowledgements

Not applicable

## Patient consent

Not applicable

## Supplementary Data

**Supplementary Fig. 1:**
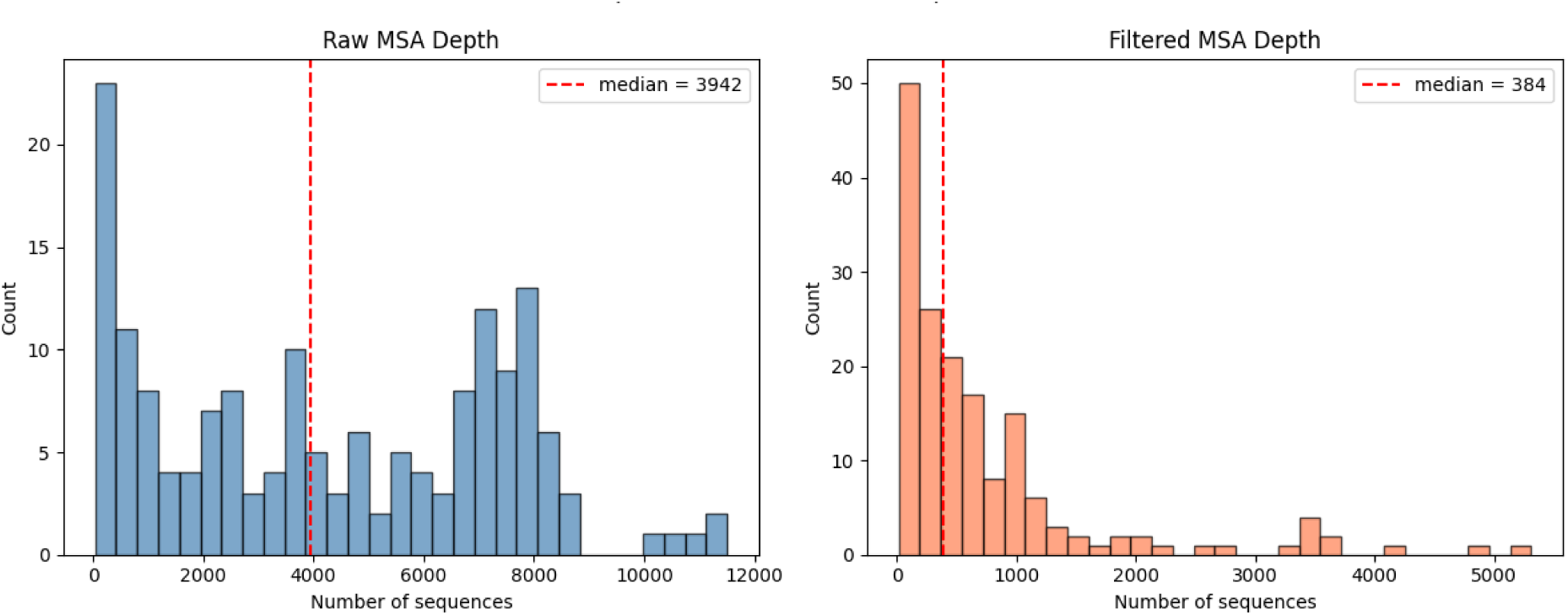
MSA distributions for original and filtered MSA following the approach of Hopf et al [43]. The original MSA distributions have a minimum depth of 34 sequences, whereas the filtered MSAs have a reduced minimum depth of 24 sequences.

**Supplementary Fig. 2:**
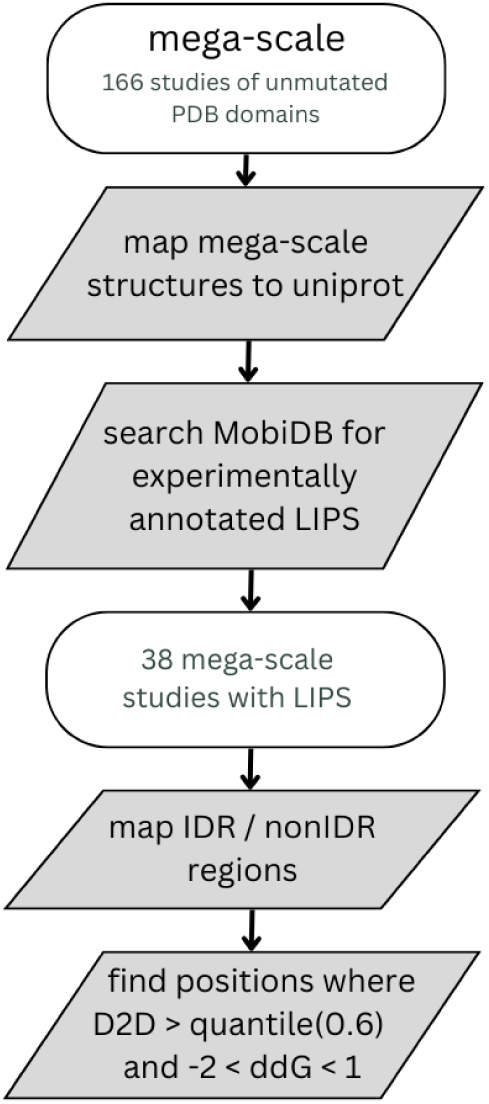
Pipeline for identifying and benchmarking LIPs using the D2D predictor. Protein structures are mapped to UniProt IDs, which are then used to retrieve functional annotations from MobiDB. For large-scale experimental datasets with available annotations, we select mutation sites that exhibit high conservation effects but low stability impacts. Predicted LIPs are compared against experimentally validated LIPs across both intrinsically disordered regions (IDRs) and structured (non-disordered) regions.

**Supplementary Fig. 3:**
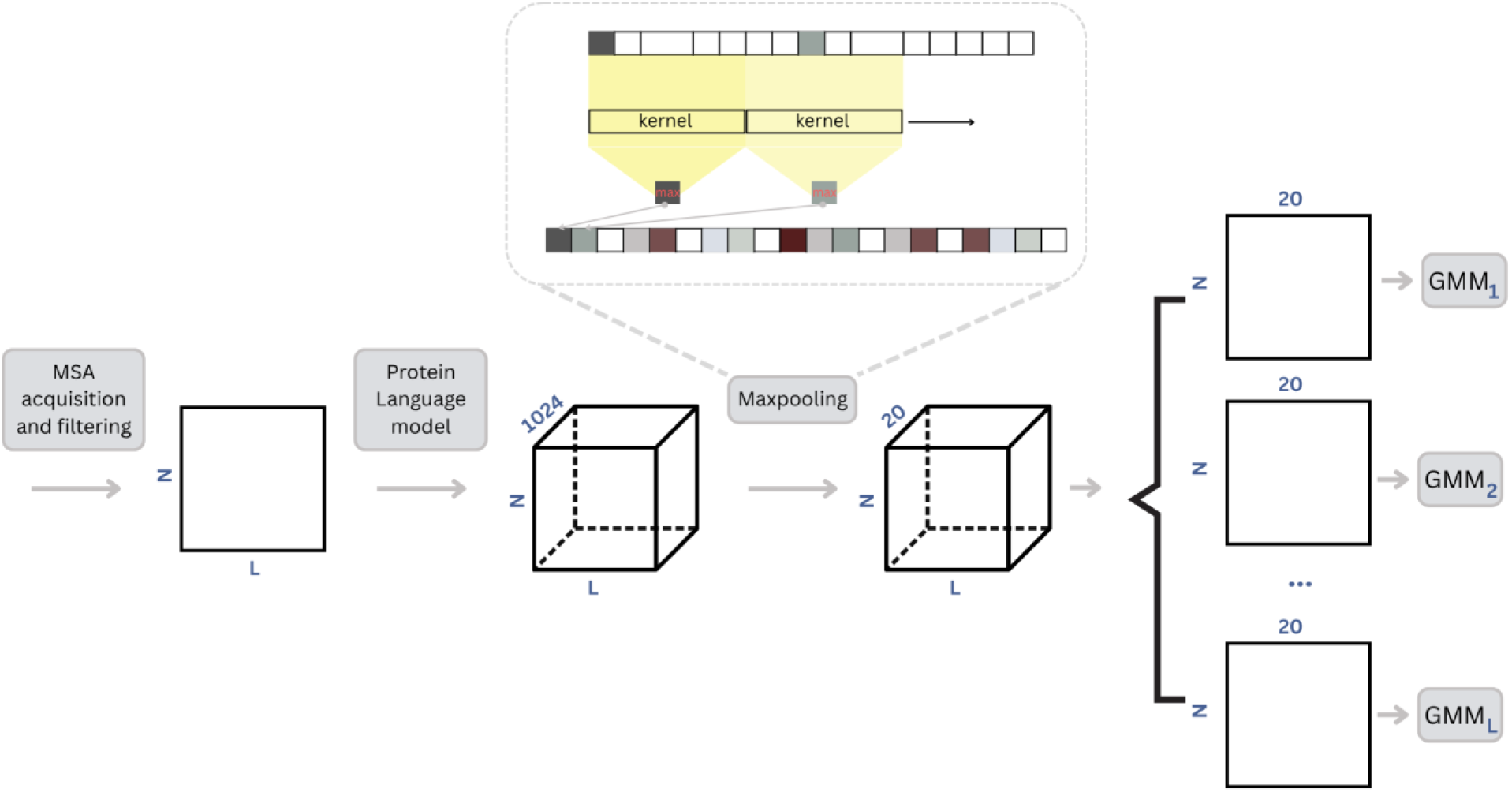
The MSA with N sequences of length L is encoded into 1024-dimensional pLM embeddings. Max pooling reduces the embedding dimensionality to 20, after which each MSA column is used as a separate input to the multivariate GMMs. The MaxPooling1D layer applies a sliding window (kernel) of fixed size across the input sequence. For each window, the maximum value is selected, reducing the sequence length and retaining key features. The stride determines how far the window moves at each step—when stride equals the kernel size, the windows do not overlap.

**Supplementary Fig. 4:**
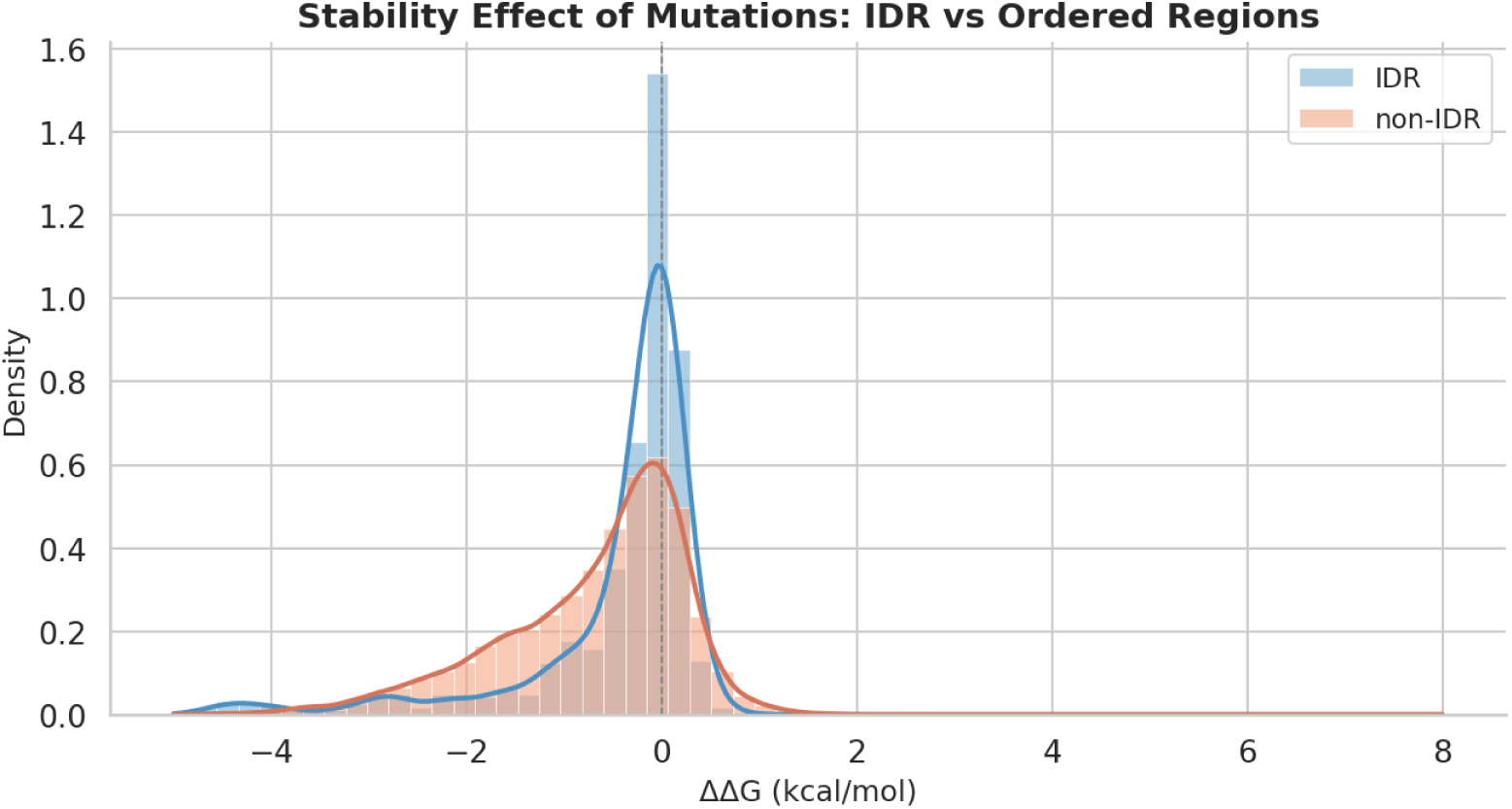
Distributions of ΔΔG values for mega-scale mutations in IDR and non-IDR regions

**Supplementary Fig. 5:**
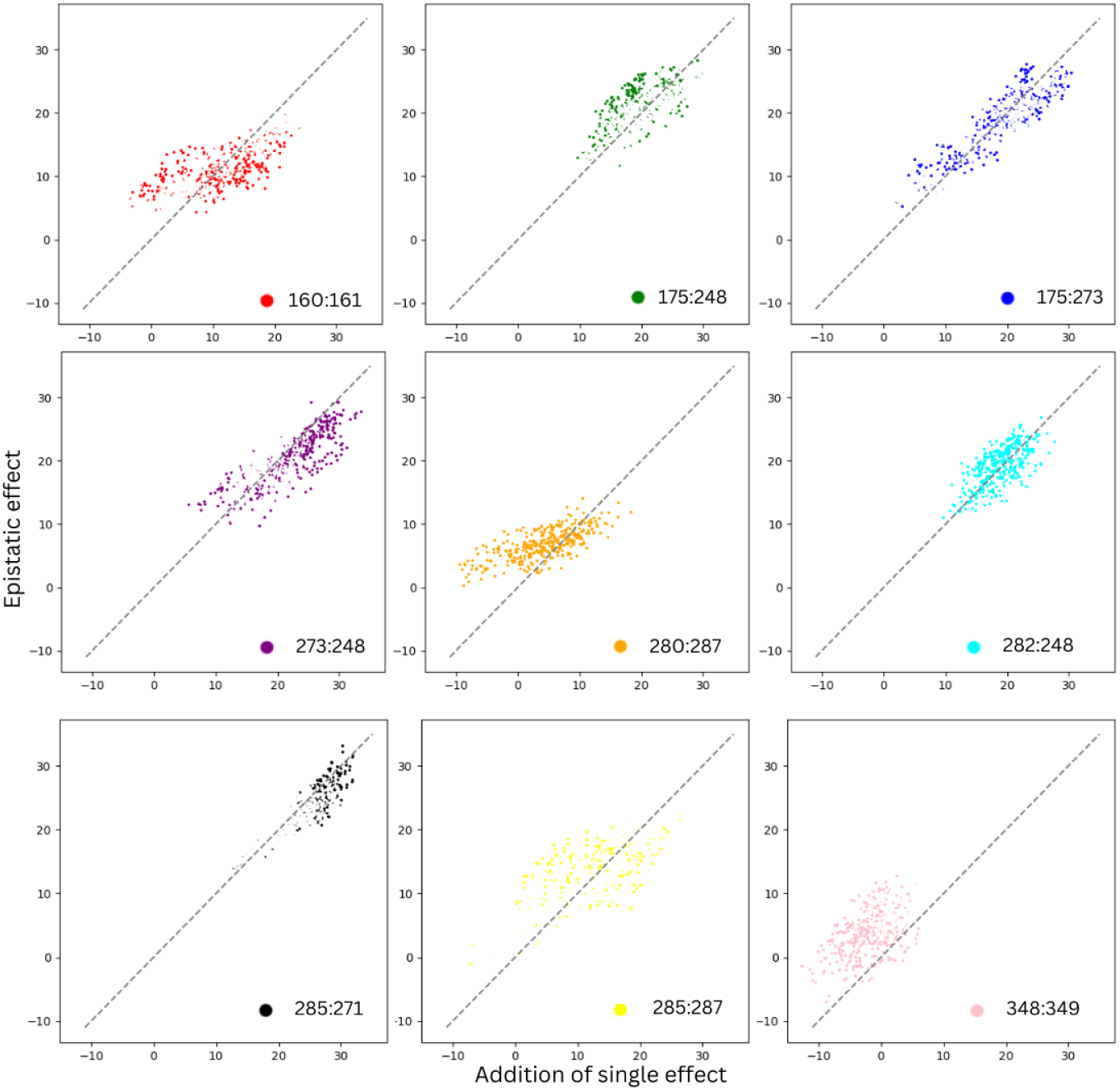
Comparison of additive vs. epistatic effects for TP53 double mutations using raw D2Deep predictions (pre-sigmoid). Mutation pairs were assessed across latent residue positions - across panels- reported by [38], evaluating all amino acid combinations. The x-axis shows the sum of individual mutation effects, while the y-axis shows the corresponding D2Deep prediction for the double mutant. This is observed, for instance, at the positions 248:249 and 285:287 that predominantly show negative epistasis effects. On the contrary, pairs such as 273:248 and 285:271 display positive epistatic interactions.

**Supplementary Table 1:** Comprehensive overview of the performance of all predictors on the CAID-3 Binding-IDR dataset.

| Predictor | AUC metric | APS metric |
| --- | --- | --- |
| bindEmbed21IDR-raw | 0.641 | 0.51 |
| bindEmbed21IDR-idr | 0.635 | 0.497 |
| **D2D** | **0.635** | **0.495** |
| LIPNet | 0.619 | 0.505 |
| ProBiPred-protein | 0.613 | 0.472 |
| AIUPred-2-binding | 0.59 | 0.463 |
| IPA-AF2-Protein-bind | 0.547 | 0.467 |
| AlphaFold-binding | 0.518 | 0.449 |
| AlphaFold3-binding | 0.511 | 0.424 |
| DRPBind-protein | 0.509 | 0.39 |
| EBIND-nucleic-acid | 0.504 | 0.251 |
| DeepDISObind-protein | 0.493 | 0.364 |

**Supplementary Table 2:**
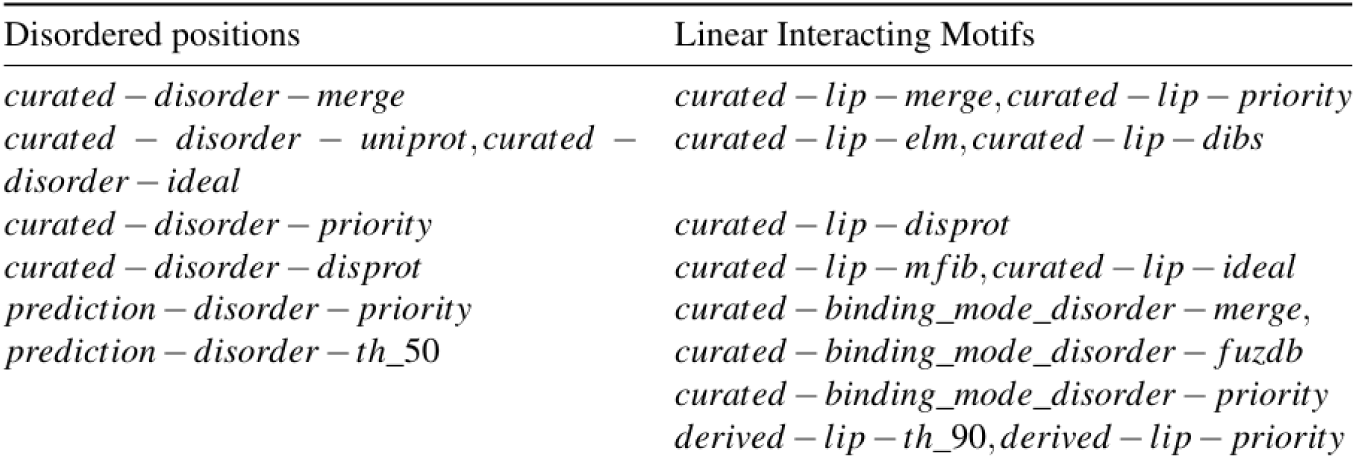
MobiDB evidence level of selected annotations.

## Notes

### Competing Interest Statement

The authors have declared no competing interest.

### Summary of Updates

Title update to 'Evolutionary constraints improve protein large language model predictions for protein stability, binding regions and epistasis'

## References

1. Peccati F, Segovia CM, Núñez-Franco R, Jiménez-Osés G. Computation of protein thermostability and epistasis.

2. National Library of Medicine (US), National Institutes of Health. Protein folding and the thermodynamic hypothesis, 1950–1962. Christian B. Anfinsen—Profiles in Science. 2019 Mar 12.

3. Surpeta B, Sequeiros-Borja CE, Brezovsky J. Dynamics, a powerful component of current and future in silico approaches for protein design and engineering. Int J Mol Sci. 2020;21(8):2713. doi:10.3390/ijms21082713.

4. Dishman AF, Volkman BF. Design and discovery of metamorphic proteins. Curr Opin Struct Biol. 2022;74:102380. doi:10.1016/j.sbi.2022.102380.

5. Domingo J, Diss G, Lehner B. Pairwise and higher order genetic interactions during the evolution of a tRNA. Nature.

6. Gonzalez CE, Ostermeier M. Pervasive pairwise intragenic epistasis among sequential mutations in TEM-1 β-lactamase. J Mol Biol. 2019;431(10):1981–1992. doi:10.1016/j.jmb.2019.03. 020.

7. Lipsh-Sokolik R, Fleishman SJ. Addressing epistasis in the design of protein function. Proc Natl Acad Sci U S A.

8. Breen MS, et al. Epistasis as the primary factor in molecular evolution. Nature. 2012;490:535–538.

9. Miton CM, Tokuriki N. How mutational epistasis impairs predictability in protein evolution and design. Protein Sci.

10. Poon A, Otto SP. Compensating for our load of mutations: freezing the meltdown of small populations. Evolution.

11. de Visser JAGM, Elena SF. The evolution of sex: empirical insights into the roles of epistasis and drift. Nat Rev Genet. 2007;8(2):139–149. doi:10.1038/nrg1985.

12. Sakai W, et al. Secondary mutations as a mechanism of cisplatin resistance in BRCA2-mutated cancers. Nature. 2008;451(7182):1116–1120. doi:10.1038/nature06633.

13. Ng PC, Henikoff S. SIFT: predicting amino acid changes that affect protein function. Nucleic Acids Res. 2003;31(13):3812–3814.

14. Adzhubei I, Jordan DM, Sunyaev SR. Predicting functional effect of human missense mutations using PolyPhen-2. Curr Protoc Hum Genet. 2013;Chapter 7:Unit7.20. doi:10.1002/0471142905.hg0720s76.

15. Raimondi D, et al. DEOGEN2: prediction and interactive visualization of single amino acid variant deleteriousness in human proteins. Nucleic Acids Res. 2017;45(W1):W201–W206. doi:10.1093/nar/gkx390.

16. Meier J, et al. Language models enable zero-shot prediction of the effects of mutations on protein function. bioRxiv. 2021. doi:10.1101/2021.07.09.450648.

17. Notin P, et al. ProteinGym: large-scale benchmarks for protein fitness prediction and design. Adv Neural Inf Process Syst. 2023;36:64331–64379.

18. Weißenow K, Heinzinger M, Rost B. Protein language-model embeddings for fast, accurate, and alignment-free protein structure prediction. Structure. 2022;30:1169–1177.

19. Ilzhöfer D, Heinzinger M, Rost B. SETH predicts nuances of residue disorder from protein embeddings. Front Bioinform. 2022;2:1019597.

20. Cui H, Wang C, Maan H, Pang K, Luo F, Duan N, Wang B. scGPT: toward building a foundation model for single-cell multi-omics using generative AI.

21. Ledzieski S, et al. Democratizing protein language models with parameter-efficient fine-tuning. Proc Natl Acad Sci U S A. 2024;121:e2405840121.

22. Rosen Y, et al. Universal cell embeddings: a foundation model for cell biology. 2023.

23. Zhang Z, Wayment-Steele HK, Brixi G, Wang H, Kern D, Ovchinnikov S. Protein language models learn evolutionary statistics of interacting sequence motifs. Proc Natl Acad Sci U S A.

24. Tzavella K, Diaz A, Olsen C, Vranken W. Combining evolution and protein language models for an interpretable cancer driver mutation prediction with D2Deep. Brief Bioinform. 2025;26(1).

25. Tsuboyama K, Dauparas J, Chen J, et al. Mega-scale experimental analysis of protein folding stability in biology and design. Nature. 2023;620:434–444.

26. Laine E, Karami Y, Carbone A. GEMME: a simple and fast global epistatic model predicting mutational effects. Mol Biol Evol. 2019;36(11):2604–2619.

27. Pires DE, Ascher DB. mCSM-AB: a web server for predicting antibody-antigen affinity changes upon mutation with graph-based signatures. Nucleic Acids Res. 2016;44(W1):W469–W473.

28. Rao RM, et al. MSA Transformer. In: Proc 38th Int Conf Mach Learn. 2021:8844–8856.

29. Meier J, et al. Language models enable zero-shot prediction of the effects of mutations on protein function. bioRxiv. 2021. doi:10.1101/2021.07.09.450648.

30. Tubiana J, Schneidman-Duhovny D, Wolfson HJ. ScanNet: an interpretable geometric deep learning model for structure-based protein binding site prediction. Nat Methods. 2022;19(6):730–739. doi:10.1038/s41592-022-01490-7.

31. Krivák R, Hoksza D. Improving protein-ligand binding site prediction accuracy by classification of inner pocket points using local features. J Cheminform. 2015;7(1):12.

32. Ramakrishnan G, Baakman C, Heijl S, et al. Understanding structure-guided variant effect predictions using 3D convolutional neural networks. Front Mol Biosci. 2023;10:1204157.

33. Pan Q, Nguyen TB, Ascher DB, Pires DEV. Systematic evaluation of computational tools to predict the effects of mutations on protein stability in the absence of experimental structures. Brief Bioinform. 2022;23(2):bbac025.

34. Caldararu O, Mehra R, Blundell TL, Kepp KP. Systematic investigation of the data set dependency of protein stability predictors. J Chem Inf Model. 2020;60(10):4772–4784.

35. Juric D, Rodon J, Tabernero J, et al. Phosphatidylinositol 3-kinase α-selective inhibition with alpelisib (BYL719) in PIK3CA-altered solid tumors: results from the first-in-human study. J Clin Oncol. 2018;36:1291–1299.

36. Juric D, Castel P, Griffith M, et al. Convergent loss of PTEN leads to clinical resistance to a PI(3)Kα inhibitor. Nature. 2015;518:240–244.

37. Vasan N, et al. Double PIK3CA mutations in cis increase oncogenicity and sensitivity to PI3Kα inhibitors. Science. 2019;366:714–723. doi:10.1126/science.aaw9032.

38. Yavuz BR, Tsai CJ, Nussinov R, et al. Pan-cancer clinical impact of latent drivers from double mutations. Commun Biol. 2023;6:202.

39. Chen S, et al. A genomic mutational constraint map using variation in 76,156 human genomes. Nature. 2024;625(7993):92–100.

40. Gonzalez CE, Ostermeier M. Pervasive pairwise intragenic epistasis among sequential mutations in TEM-1-lactamase. J Mol Biol. 2019;431(10):1981–1992. doi:10.1016/j.jmb.2019.03.020.

41. Sakai W, et al. Secondary mutations as a mechanism of cisplatin resistance in BRCA2-mutated cancers. Nature. 2008;451:1116.

42. Kamaraj B, Bogaerts A. Structure and function of p53-DNA complexes with inactivation and rescue mutations: a molecular dynamics simulation study. PLoS One. 2015;10(8):e0134638. doi:10.1371/journal.pone.0134638.

43. Hopf TA, Ingraham JB, Poelwijk FJ, et al. Mutation effects predicted from sequence co-variation. Nat Biotechnol. 2017;35:128–135. doi:10.1038/nbt.3769.

